# PlasmidFlow: A Web-based Platform for Interactive Visualization of Plasmid-Driven Traits

**DOI:** 10.1101/2025.08.17.670741

**Authors:** Adeel Farooq, Asma Rafique

**Affiliations:** Canadian Research Institute for Food Safety, Department of Food Science, University of Guelph, Guelph N1G 2W1, Canada; Department of Human Health Sciences, University of Guelph, Guelph N1G 2W1, Canada

**Keywords:** Plasmids, Horizontal Gene Transfer, Flow Diagram, Trait Network Graph, Microbial Genomics, Interactive Visualization, Antibiotic Resistance

## Abstract

Plasmids are central mediators of horizontal gene transfer (HGT), facilitating the spread of antimicrobial resistance, virulence determinants, and adaptive functions across microbial communities. Despite the availability of extensive plasmid sequencing data, visualization of plasmid-mediated traits remains hindered by reliance on static figures or coding-intensive pipelines, limiting accessibility for microbiologists and epidemiologists. We present **PlasmidFlow** (https://plasmidflow.metabopotential.site), an interactive, web-based platform for the exploration of plasmid–host–trait associations across ecological and clinical contexts. The system integrates structured data parsing, relational storage, and modular visualization to support real-time exploration without requiring programming expertise. Users can upload plasmid–host–trait datasets, apply environment-or trait-specific filters, and generate publication-ready Sankey diagrams, plasmid-sharing network graphs, and binary trait heatmaps with full customization of visual properties. High-resolution export in SVG, PDF, and PNG formats, along with reproducible session storage, ensures compatibility with both research and publication workflows. By combining interactivity, scalability, and reproducibility, PlasmidFlow addresses a critical gap in microbial genomics, providing a user-friendly and extensible resource for investigating plasmid-driven traits in the context of antimicrobial resistance surveillance, microbial ecology, and evolutionary biology.

## Introduction

Plasmids are extrachromosomal, self-replicating DNA elements that play a pivotal role in microbial adaptability and evolution. Through horizontal gene transfer (HGT), they act as vehicles for disseminating genes associated with antibiotic resistance, virulence, and metabolic innovation, thereby shaping microbial ecology and clinical outcomes (Frost et al., 2005; Smillie et al., 2010; Norman et al., 2009). The global rise of antimicrobial resistance (AMR) is tightly linked to plasmid-mediated mobility of resistance determinants, highlighting the urgent need for systematic tools to study plasmid distributions and their encoded traits (Carattoli, 2013; Partridge et al., 2018). In addition to AMR, plasmids contribute to the spread of toxin–antitoxin systems, secretion systems, and mobile genetic elements that enhance niche colonization and survival (Hall et al., 2017; Arredondo-Alonso et al., 2020).

Advances in sequencing technologies have generated vast repositories of plasmid data across clinical, environmental, and metagenomic contexts (Krawczyk et al., 2018; Zhou & Olman, 2007). However, translating these datasets into biological insight remains challenging. Conventional bioinformatics pipelines rely on static figures or coding-intensive scripts, which can limit accessibility for microbiologists, clinicians, and public health researchers who may not have computational expertise. Moreover, existing visualization approaches often lack interactive features for filtering, customizing, or comparing plasmid-mediated traits across ecological niches (Huse et al., 2014; Ondov et al., 2019).

To address these challenges, we developed PlasmidFlow, an interactive, browser-based platform that integrates dynamic data handling, trait-based filtering, and advanced visualization to facilitate exploration of plasmid-driven traits. Unlike static visualization methods, PlasmidFlow enables real-time generation and customization of Sankey diagrams, network graphs, and trait heatmaps, all exportable in publication-ready formats. The platform was designed to be both user-friendly and extensible, requiring no programming expertise while remaining scalable for large datasets. By bridging the gap between raw plasmid sequence data and intuitive visualization, PlasmidFlow supports hypothesis generation in microbial ecology, epidemiology, and evolutionary biology, with particular relevance to antimicrobial resistance surveillance and microbial trait discovery.

### System Architecture and Interface Design

PlasmidFlow adopts a modular client–server architecture designed for portability, scalability, and reproducibility. The client side, accessible through any modern web browser, is organized into three main panels: a navigation header and sidebar for layout and branding, a data panel for uploading plasmid–host–trait datasets and applying environmental or trait-based filters, and a visualization panel for customizing graph properties and previewing results interactively. On the server side, uploaded datasets are validated, transformed, and stored in a relational backend. By default, PlasmidFlow uses SQLite, an embedded SQL database engine optimized for lightweight persistence (Hipp, 2022), while PostgreSQL provides support for scalability and concurrent querying in larger deployments (Stonebraker & Rowe, 1986). This modular separation of interface, processing, and storage ensures that PlasmidFlow remains adaptable across personal workstations, institutional servers, and cloud environments (**Figure 1)**.

**Figure 1.**
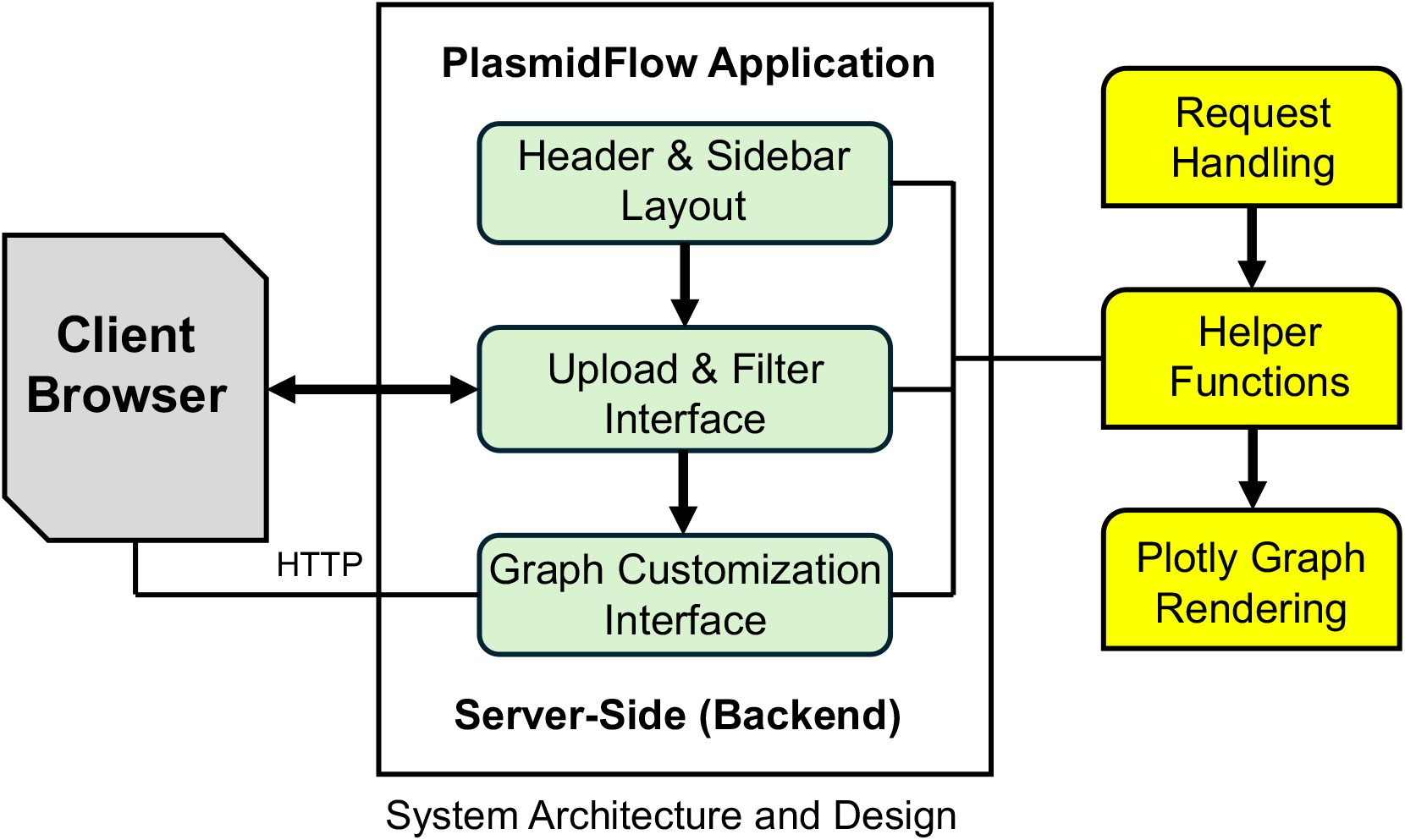
System architecture of the PlasmidFlow web interface. The client-side browser interacts with a modular Dash application composed of three main interface layers: (1) the header and sidebar layout, (2) the upload and filtering interface, and (3) the graph customization panel. These frontend components communicate with backend modules responsible for request handling, data processing through helper functions, and dynamic rendering of visualizations using Plotly.

### Methods and Implementation

The front end of PlasmidFlow is developed using HTML5 for structural markup (W3C, 2014), Bootstrap for responsive design (Otto & Thornton, 2011), and JavaScript for dynamic browser interactivity (ECMA International, 2015). Dynamic components are served through Flask, a Python web micro-framework for routing and session management (Grinberg, 2018), which communicates with the client using AJAX-based asynchronous requests (Garrett, 2005). Users can upload CSV-formatted datasets, select traits of interest, and instantly preview processed data through a responsive interface.

On the backend, data parsing, validation, and transformation are handled using pandas (McKinney, 2011) and NumPy (Harris et al., 2020), ensuring efficient manipulation of tabular and numerical data. Visualizations are dynamically generated using Plotly for interactive charting (Plotly Technologies Inc., 2015) and NetworkX for network-based analysis (Hagberg, Schult, & Swart, 2008). Sankey diagrams represent directional plasmid-mediated flow, trait-sharing networks illustrate ecological overlap, and binary or frequency-scaled heatmaps display trait distributions. A dedicated customization panel enables users to modify visualization attributes such as color scales, node and edge properties, and font sizes, with settings serialized in JSON format to support reproducibility **(Figure 2)**.

**Figure 2.**
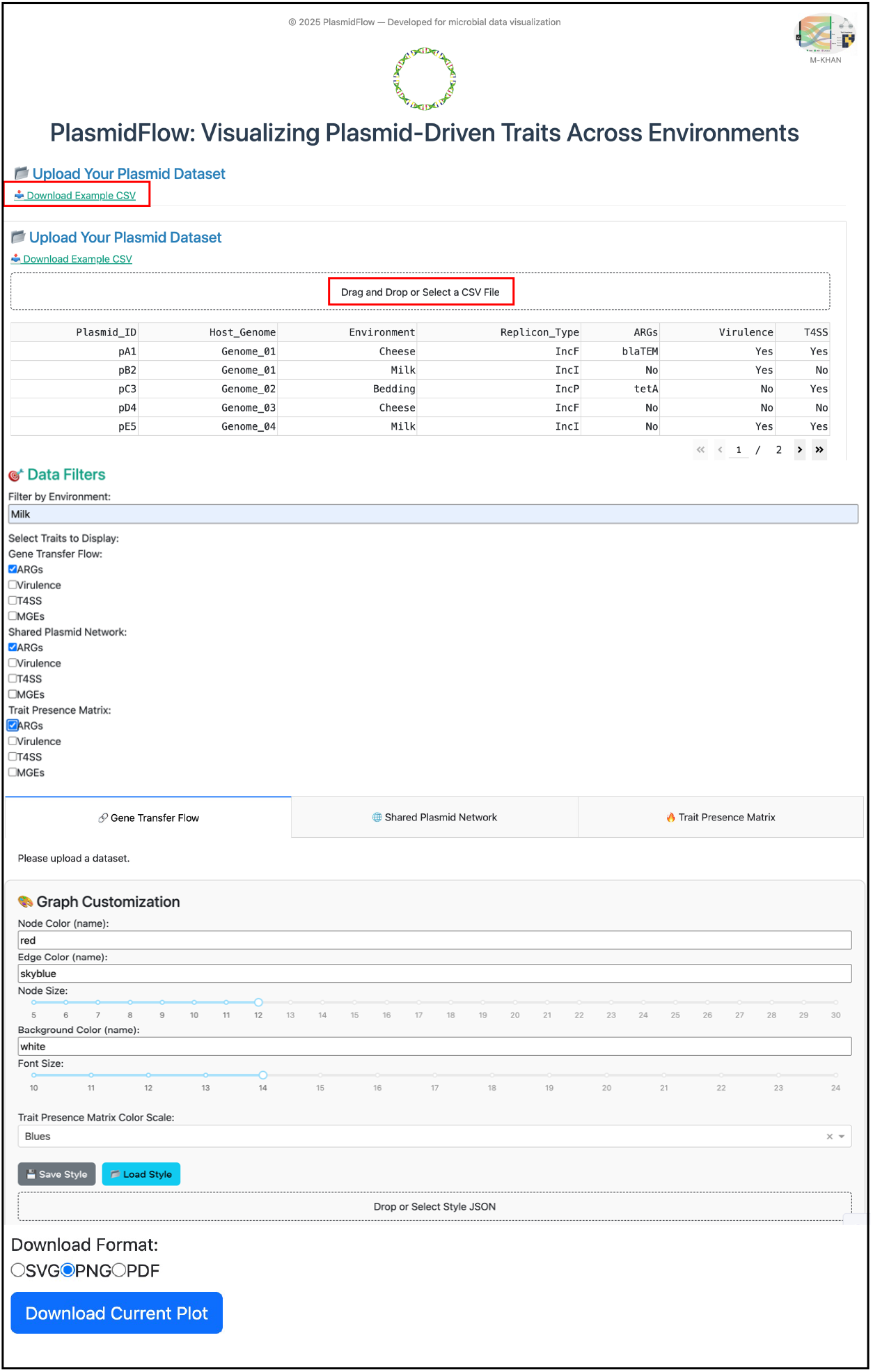
Interactive dashboard layout of the PlasmidFlow web application. The interface consists of modular panels for uploading plasmid datasets, selecting environments and traits, and customizing visual outputs. Users can drag and drop CSV files containing plasmid-host-trait metadata and filter the dataset by environment or gene transfer feature (e.g., ARGs, virulence factors, T4SS, MGEs). Visualization modes include Sankey diagrams, network graphs, and trait heatmaps. A customization panel allows real-time adjustments to graph appearance, including node color, edge color, font size, and background. Export options are provided in SVG, PNG, and PDF formats for publication-ready figures.

To ensure dissemination and publication-ready outputs, PlasmidFlow integrates high-resolution export of figures in SVG, PNG, and PDF formats using the Kaleido engine (Sievert, 2020). Combined with session persistence and database storage, this pipeline guarantees that analyses are transparent, reproducible, and accessible to both coding and non-coding users.

### Use Case and Results

To demonstrate the functionality of PlasmidFlow, we applied the platform to a synthetic dataset modeling plasmid–host–trait associations across diverse environments, including soil, river water, marine ecosystems, and clinical isolates. The dataset contained plasmid identifiers, host bacterial genomes, environmental metadata, replication types, and annotated traits such as antimicrobial resistance genes (ARGs), virulence determinants, type IV secretion systems (T4SS), and mobile genetic elements (MGEs). This dataset is bundled within the application and available to all users through the “Download Example CSV” feature, ensuring reproducibility and enabling first-time users to explore PlasmidFlow without the need for custom inputs.

Upon uploading the dataset, PlasmidFlow immediately parsed and displayed a structured preview of plasmid–trait associations. Users were able to filter by environment (e.g., clinical vs. marine) or by trait category (e.g., ARGs vs. virulence factors), enabling rapid subsetting of the data. The interactive visualization modules revealed distinct insights across the three modes of display. Sankey diagrams **(Figure 3, top)** illustrated directional plasmid-mediated flow, with wide connections between clinical isolates and ARG-associated plasmids, highlighting their disproportionate contribution to antimicrobial resistance dissemination (Partridge et al., 2018). In contrast, plasmids in soil and aquatic environments showed greater trait diversity, including virulence factors and MGEs, consistent with ecological heterogeneity (Hall et al., 2017).

**Figure 3.**
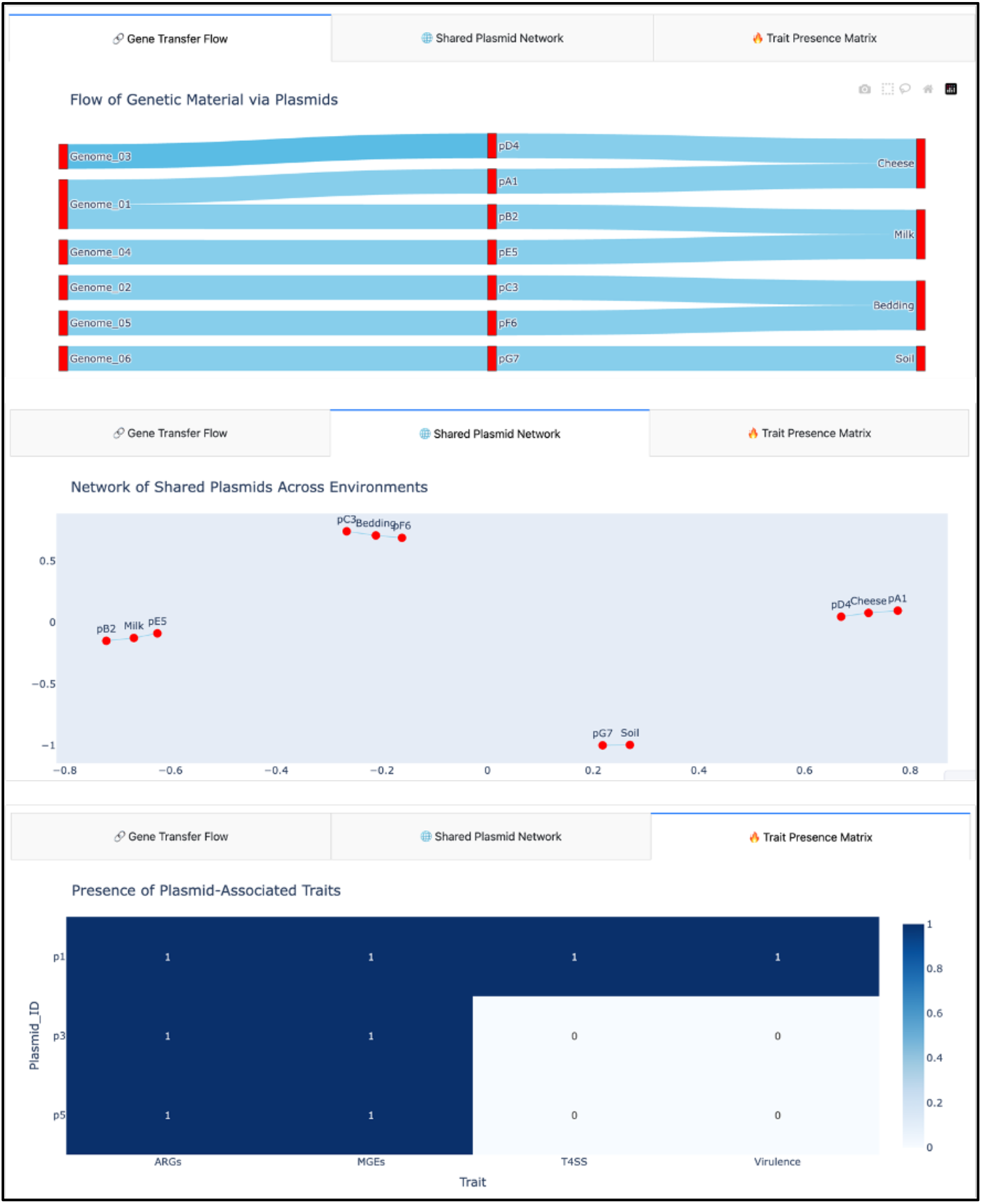
Visualization outputs generated by PlasmidFlow for plasmid-mediated gene transfer and trait distribution. Top: Sankey diagram illustrating the directional flow of plasmid-mediated genetic material from bacterial genomes to environmental compartments. Each link represents a plasmid transfer, with node width proportional to the number of associations. Middle: Shared plasmid network showing ecological connectivity among environments based on plasmid overlap. Nodes represent plasmids or environments, and edges indicate shared elements. Bottom: Trait presence matrix displaying the distribution of plasmid-associated features (e.g., ARGs, T4SS, MGEs, virulence factors) across selected plasmids. Each cell is color-coded to indicate binary presence (1) or absence (0) of a trait.

The plasmid-sharing network graph **(Figure 3, middle)** highlighted clusters of plasmids bridging environmental and clinical compartments. Network centrality analysis identified a subset of plasmids that were shared across multiple environments, suggesting ecological connectivity and the potential for cross-compartmental transfer events (Norman et al., 2009; Arredondo-Alonso et al., 2020). Visualization controls enabled users to customize node size by plasmid abundance and edge thickness by co-occurrence frequency, facilitating detection of ecological hubs and hotspots of plasmid-mediated exchange.

Finally, the trait heatmap (**Figure 3, bottom)** provided a binary presence–absence matrix of traits across plasmids. This mode revealed strong enrichment of ARGs in clinical plasmids and broader distributions of secretion systems and MGEs in environmental datasets. By adjusting color scales and cluster ordering, users could quickly compare trait repertoires between environments. All visualizations were exportable in high resolution (SVG, PNG, PDF), ensuring suitability for scientific publication and downstream analysis.

Collectively, these results demonstrate that PlasmidFlow provides an intuitive, flexible, and reproducible environment for visualizing plasmid-driven traits. The case study illustrates how the platform can be used to identify ecological connectivity, trait clustering, and differential enrichment of plasmid-borne functions across microbial niches.

## Availability and Future Directions

PlasmidFlow is open-source and available at https://plasmidflow.metabopotential.site.

Planned enhancements include support for expanded trait ontologies, multi-sample dashboards, genome browser modules, and session bookmarking for reproducibility.

## Author Contributions

Conceptualization, A.F.; methodology, A.F. and A.R.; software, A.F.; validation, A.F., and A.R.; formal analysis, A.F.; investigation, A.F.; resources, A.F.; data curation, A.F.; writing—original draft preparation, A.F. and A.R.; writing—review and editing, A.F.; visualization, A.F.;. All authors have read and agreed to the published version of the manuscript.

## Data Availability Statement

The data supporting the findings of this study are available from the corresponding author upon reasonable request.

## Acknowledgments

This work was developed independently and is dedicated to the memory of M. Khan.

## References

Arredondo-Alonso, S., Willems, R. J. L., van Schaik, W., & Schürch, A. C. (2020). On the (im)possibility of reconstructing plasmids from whole-genome short-read sequencing data. Microbial Genomics, 6(10), e000384.

Carattoli, A. (2013). Plasmids and the spread of resistance. International Journal of Medical Microbiology, 303(6–7), 298–304.

ECMA International. (2015). ECMAScript® 2015 Language Specification (6th Edition). Geneva, Switzerland.

Frost, L. S., Leplae, R., Summers, A. O., & Toussaint, A. (2005). Mobile genetic elements: the agents of open-source evolution. Nature Reviews Microbiology, 3(9), 722–732.

Garrett, J. J. (2005). Ajax: A new approach to web applications. Adaptive Path. Retrieved from https://adaptivepath.org/ideas/ajax-new-approach-web-applications/

Grinberg, M. (2018). Flask Web Development: Developing Web Applications with Python. O’Reilly Media.

Hagberg, A. A., Schult, D. A., & Swart, P. J. (2008). Exploring network structure, dynamics, and function using NetworkX. In Proceedings of the 7th Python in Science Conference (pp. 11–15).

Hall, J. P. J., Brockhurst, M. A., & Harrison, E. (2017). Sampling the mobile gene pool: innovation via horizontal gene transfer in bacteria. Philosophical Transactions of the Royal Society B, 372(1735), 20160424.

Harris, C. R., Millman, K. J., van der Walt, S. J., Gommers, R., Virtanen, P., Cournapeau, D., Oliphant, T. E. (2020). Array programming with NumPy. Nature, 585, 357–362.

Hipp, D. R. (2022). SQLite. Retrieved from https://www.sqlite.org

Huse, S. M., Mark Welch, D. B., Voorhis, A., Shipunova, A., Morrison, H. G., Eren, A. M., & Sogin, M. L. (2014). VAMPS: a website for visualization and analysis of microbial population structures. BMC Bioinformatics, 15, 41.

Krawczyk, P. S., Lipinski, L., & Dziembowski, A. (2018). PlasFlow: predicting plasmid sequences in metagenomic data using genome signatures. Nucleic Acids Research, 46(6), e35.

McKinney, W. (2011). pandas: a foundational Python library for data analysis. In Proceedings of the 9th Python in Science Conference (pp. 51–56).

Norman, A., Hansen, L. H., & Sørensen, S. J. (2009). Conjugative plasmids: vessels of the communal gene pool. Philosophical Transactions of the Royal Society B, 364(1527), 2275–2289.

Ondov, B. D., Treangen, T. J., Melsted, P., Mallonee, A. B., Bergman, N. H., Koren, S., & Phillippy, A. M. (2019). Mash: fast genome and metagenome distance estimation using MinHash. Genome Biology, 16, 132.

Otto, M., & Thornton, J. (2011). Bootstrap. Retrieved from https://getbootstrap.com

Partridge, S. R., Kwong, S. M., Firth, N., & Jensen, S. O. (2018). Mobile genetic elements associated with antimicrobial resistance. Clinical Microbiology Reviews, 31(4), e00088–17.

Plotly Technologies Inc. (2015). Collaborative data science. Montréal, QC. Retrieved from https://plot.ly

Sievert, C. (2020). Interactive Web-Based Data Visualization with R, plotly, and shiny. Chapman & Hall/CRC.

Smillie, C., Garcillán-Barcia, M. P., Francia, M. V., Rocha, E. P. C., & de la Cruz, F. (2010). Mobility of plasmids. Microbiology and Molecular Biology Reviews, 74(3), 434–452.

Stonebraker, M., & Rowe, L. A. (1986). The design of POSTGRES. In Proceedings of the 1986 ACM SIGMOD International Conference on Management of Data (pp. 340–355). ACM.

W3C. (2014). HTML5: A vocabulary and associated APIs for HTML and XHTML. W3C Recommendation. Retrieved from https://www.w3.org/TR/html5

Zhou, F., & Olman, V. (2007). Visualization of metagenomic data. Briefings in Bioinformatics, 8(6), 526–545.

